# Genome sequence of the wheat stem sawfly, *Cephus cinctus*, a primitive hymenopteran and wheat pest, illuminates evolution of hymenopteran chemoreceptors

**DOI:** 10.1101/380873

**Authors:** Hugh M. Robertson, Robert M. Waterhouse, Kimberly K. O. Walden, Livio Ruzzante, Maarten J. M. F. Reijnders, Brad S. Coates, Fabrice Legeai, Joanna C. Gress, Sezgi Biyiklioglu, David K. Weaver, Kevin W. Wanner, Hikmet Budak

## Abstract

The wheat stem sawfly, *Cephus cinctus*, is a major pest of wheat and key ecological player in the grasslands of western North America. It also represents a distinctive lineage of sawflies that appeared early during the hymenopteran radiation, but after the clade of Eusymphyta sawflies that is the sister lineage of all other Hymenoptera. We present a high-quality draft genome assembly of 162 Mbp in 1,976 scaffolds with a scaffold N50 of 622 kbp. Automated gene annotation identified 11,210 protein-coding gene models and 1,307 non-coding RNA models. Thirteen percent of the assembly consists of ~58,000 transposable elements partitioned equally between Class-I and Class-II elements. Orthology analysis reveals that 86% of *Cephus* proteins have identifiable orthologs in other insects. Phylogenomic analysis of conserved subsets of these proteins supports the placement of the Cephidae between the Eusymphyta and the parasitic woodwasp superfamily Orussoidea. Manual annotation and phylogenetic analysis of families of odorant, gustatory, and ionotropic receptors, plus odorant binding proteins, shows that *Cephus* has representatives for most conserved and expanded gene lineages in the Apocrita (wasps, ants, and bees). *Cephus* has also maintained several insect gene lineages that have been lost from the Apocrita, most prominently the carbon dioxide receptor subfamily. Furthermore, *Cephus* encodes a few small lineage-specific chemoreceptor gene family expansions that might be involved in adaptations to new grasses including wheat. These comparative analyses identify gene family members likely to have been present in the hymenopteran ancestor and provide a new perspective on the evolution of the chemosensory gene repertoire.

## Introduction

The wheat stem sawfly, *Cephus cinctus* Norton, is a major pest of wheat in western North America with a southward expanding geographic range likely driven by localized adaptation to cultivated crops from surrounding wildlands (Beres et al. 2011; Lesieur et al. 2016; Adhikari et al. 2018; Varella et al. 2018). It is a native species that also uses many other grass hosts and hence plays an important role in the ecology of grasslands (Cockrell et al. 2017). Worldwide, as insect burdens on wheat farmers, members of the genus *Cephus* rival the Hessian fly, *Mayetiola destructor*, a dipteran for which a genome sequence is available (Zhao et al. 2015). Female sawflies lay eggs in growing wheat stems and the larvae grow within the stem lumen (Figure 1). Once in the final instar, the sole survivor of larval cannibalism in each stem retreats to the base of the stem, which it girdles in preparation for pupation (Buteler et al. 2009; 2015). Economic loss is from both the larval presence and the subsequent collapse or lodging of the wheat stem (Bekkerman and Weaver 2018). As a result of this life history, wheat stem sawflies are protected from control using conventional sprayed insecticides. The development of alternative approaches relies on an improved understanding of *C. cinctus* chemical ecology (Cossé et al. 2002; Weaver et al. 2009) and associated behaviors on host plants (Buteler et al. 2009; Varella et al. 2017), but large-scale molecular biology studies to date remain limited to antennal transcriptomics that examined chemosensory genes (Gress et al. 2013) and analyses of larval and adult RNA sequencing (RNA-seq) data that focused on the identification of non-coding RNA transcripts (Cagirici et al. 2017). A key first step in this direction is to build genomic and additional transcriptomic resources for this sawfly that could, for example, facilitate the development of new resistant wheat strains using expression of RNA interference (RNAi) constructs against essential wheat stem sawfly genes.

**Fig. 1.**
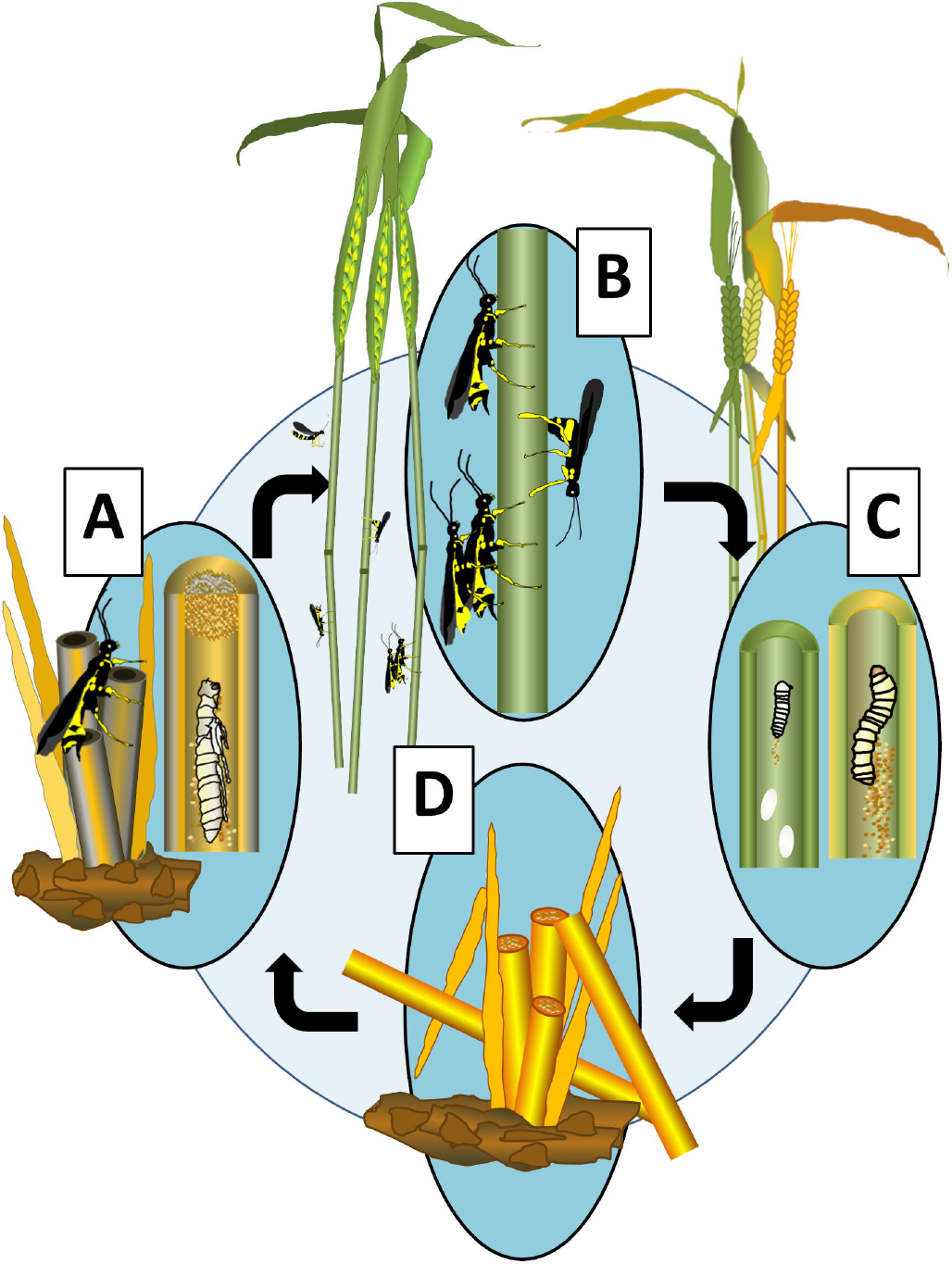
Life cycle of the wheat stem sawfly, *Cephus cinctus* Norton. A. After overwintering in diapause in a senesced wheat stem, a pre-pupa metamorphoses in late spring or early summer. The pupal stage lasts 10-12 days, after which the newly-eclosed adult chews a hole in a frass plug left by the larva to emerge from the stem. B. The adults can mate immediately after emergence, either on the wheat stubble or on new host stems. The short-lived females oviposit 30-40 eggs before dying in 7-10 days. C. Newly-deposited eggs in the stem lumen, usually from multiple females, hatch in 5-10 days. The neonate larvae feed on the parenchyma lining the stem interior. As the larvae grow through 4-5 larval instars they forage on the stem lining throughout the stem by boring through the nodes. The duration of larval feeding can range from 3-8 weeks. D. Within the stem, cannibalism leads to a single surviving larva that migrates to the base of the stem at plant senescence and ripening. The larva girdles the interior of the ripened stem wall, cutting a groove that encircles the entire stem. Wind or gravity causes the weakened stem to break and lodge, leaving the unharvested wheat head on the soil. Below this groove, the larva makes a frass plug before migrating to the base of the stem near the crown. Within a thin hibernaculum, the pre-pupa remains protected in obligate diapause from late summer through to warming the following spring. This period of inactivity can last more than 9 months. *Image created by Megan L. Hofland and Norma J. Irish, Department of Land Resources and Environmental Sciences, Montana State University*

An additional significance to this insect is that the genus *Cephus* represents the sawfly family Cephidae and superfamily Cephoidea, and molecular phylogenomic analysis of the Hymenoptera suggests that this is a distinctive lineage of sawflies that radiated after the more well-known and basal sawflies such as the family Tenthredinidae, now called the Eusymphyta, and before the parasitic woodwasp superfamily of Orussoidea (Peters et al. 2017). It therefore represents an interim step in the origin of the major lineage of the Apocrita, and so provides an important comparator and outgroup for understanding some of the peculiarities of the genomes, gene contents, and biology of the wasps, ants, and bees. Genomic resources for *C. cinctus* are therefore not only important for pest control but also contribute to augmenting the growing representation of sequenced hymenopteran genomes (Branstetter et al. 2018).

Here we report a draft genome assembly and annotation for *C. cinctus*, with genome-wide analyses of protein orthology and repeat content, and phylogenomic analysis that supports the placement of Cephoidea between Eusymphyta and Orussoidea. Supported by extensive manual gene annotation efforts, our phylogenetic analyses focused on four major gene families encoding chemosensory proteins of ecological importance. *Cephus*, as a member of an early radiating lineage, provides a new perspective on the likely ancestral repertoires and subsequent lineage-specific evolution of odorant binding proteins (OBPs), and odorant, gustatory, and ionotropic receptors (ORs, GRs, and IRs), across the Hymenoptera.

## Materials and Methods

### DNA isolation, genomic library preparation, sequencing, and genome assembly

Genomic DNA was isolated from a single male and separately from pooled females (Table S1). Briefly, insects were ground in liquid nitrogen before lysing in a SDS solution overnight with Proteinase K. The homogenate was treated with RNaseA, and proteins/debris were collected after high-salt precipitation and centrifugation. After ethanol precipitation, the DNA was resuspended in 10 mM Tris and evaluated on an agarose gel and by Qubit quantification. The following libraries were generated for sequencing: 500 bp and 1.5 kb insert shotgun libraries from the single male sawfly, and 3 and 5 kb insert mate-pair libraries from the pooled female DNA. The 500 bp and 1.5 kb insert shotgun libraries were prepared with Illumina TruSeq DNAseq Sample Prep kit. The 3 kb and 5 kb mate-pair libraries were prepared similarly except a custom linker was ligated between the fragment ends to facilitate mate-pair recovery. All libraries were sequenced for at least 100 cycles on Illumina GAIIx or HiSeq2000 machines using Illumina TruSeq SBS Sequencing kit v3. Bases were called with Casava v1.6 or 1.82. The custom 3 kb and 5 kb mate-pair libraries were filtered for properly-oriented reads of the appropriate insert size and uniqueness using in-house custom pipeline scripts. Raw Illumina reads were 5′- and 3′-trimmed for nucleotide-bias and low-quality bases using the FASTX Toolkit (http://hannonlab.cshl.edu/fastx_tookit/). Trimmed reads were error-corrected by library with Quake (Kelley et al. 2010) counting 19-mers. SOAPdenovo v2.04 (Luo et al. 2012) was employed with K=49 to assemble the 500 bp-insert shotgun library reads followed by scaffolding with iteratively longer-insert shotgun and mate-pair libraries and use of GapCloser v1.12 to close gaps generated in the scaffolding (Luo et al. 2012).

### RNA isolation, RNAseq library preparation, sequencing, and transcriptome assembly

We undertook RNAseq studies of whole animals and tissues relevant to chemosensation such as antennae, heads without antennae, abdomen tips, and abdomens without tips (Table S1). Entire animals or body parts were ground in 1ml Trizol in glass tissue grinders and filtered over a Qiagen Qiashredder column. The homogenate was extracted with chloroform and the RNA was precipitated with linear polyacrylamide (10mg/mL) and isopropanol. RNA pellets were washed with 75% ethanol and resuspended in RNase-free water. RNA was quantified with a Qubit RNA Broad Range Assay Kit on a Qubit fluorometer (Life Technologies). RNA was visualized using ethidium bromide on a 1.0% agarose gel or on a BioAnalyzer (Agilent Genomics). The RNAseq libraries were prepared from an average cDNA fragment size of 250 bases using the Illumina TruSeq Stranded RNAseq Sample Prep kits. The libraries were individually barcoded and quantitated using qPCR before pooling and sequencing from both ends with TruSeq SBS Sequencing kits v2 or 3 for 100 cycles on a HiSeq2000 or with a HiSeq4000 Sequencing kit v1 for 150 cycles on a HiSeq4000 instrument. Bases were called with Casava v1.6 or 1.82 or bcl2fastq v2.17.1.14. Reads were trimmed as for DNA sequencing. The resulting trimmed reads from the male and female 2011 and the male and female antennal 2013 samples were assembled using Trinity (Release 2014-04-13) (http://trinityrnaseq.github.io/) (Haas et al. 2013).

### Automated gene modeling

Annotation of the Cephus genome assembly was performed by the NCBI using their Eukaryotic Genome Annotation Pipeline (https://www.ncbi.nlm.nih.gov/genome/annotation_euk/process/), with experimental support from the RNAseq and transcriptome. The results are available at www.ncbi.nlm.nih.gov/genome/annotation_euk/Cephus_cinctus/101/, and constitute Ccin0GSv1.0 (Official Gene Set).

### Manual gene modeling

Full descriptions of gene modeling methods are presented in the Online Supplement for each gene family. Briefly, TBLASTN searches of the genome assembly were performed using relevant proteins from other Hymenoptera and other insects, at stringencies relevant to the various families, at the i5k Workspace@NAL (Poelchau et al. 2014) where the genome assembly is presented. Relevant NCBI gene models were examined in the linked browser and modified if necessary in light of gene model information from other species, as well as the extensive RNAseq, which was mapped to the genome and informed exon-intron boundaries. The 5’ and 3’ UTRs of the automated models were carefully adjusted based on the RNAseq mapping, and any extensive overlaps with the 5’ or 3’ UTRs of highly-expressed neighboring genes were noted to avoid problems with subsequent analysis of expression levels. Genes were named in the Apollo browser according to relevant approaches for each gene family (Supplemental material), and the manually modeled and annotated genes were merged with the NCBI models to generate CcinOGSv1.1, which is available from i5k Workspace@NAL. All new protein sequences are available in FASTA format in Auxiliary file 3. Phylogenetic methods are detailed in the supplement and alignments are available from HMR.

### microRNA Annotation

microRNA annotation was performed as outlined previously (Cagrici et al., 2017). High-confidence microRNAs for Hexapoda were downloaded from miRBase.org (Release 21) and a query was formed for a homology-based screen of the genome assembly.

### Phylogenomics, orthology, gene set and genome completeness

The maximum likelihood molecular species phylogeny (Figure 2) was estimated from the concatenated protein sequence superalignment of 852 single-copy orthologs, using the automated gene models of CcinOGSv1.0 extracted from OrthoDB v9.1 (Zdobnov et al. 2017), using the 17 species shown in Table S3. The presence, absence, and copy-numbers of orthologs from 116 insect species were assessed to partition genes from each species into the distribution of categories shown in the bar chart (Figure 2). For each of the two sawflies and the woodwasp, percent amino acid identities with orthologs from *Nasonia vitripennis, Pogonomyrmex barbatus*, and *Apis mellifera*, were calculated from the trimmed multiple sequence alignments of 1,035 single-copy Hymenoptera orthologs (Figure 2, violin plots). Gene Ontology (GO) terms from *Drosophila melanogaster, A. mellifera, Tribolium castaneum*, and *Anopheles gambiae* were compared to identify *C. cinctus* genes with orthologs associated with GO terms assigned to at least two (i.e. annotations in common), just one (i.e. unique annotations), or none of the four species. Completeness in terms of expected gene content for the genomes and annotated gene sets of each of the eight analysed hymenopteran species was assessed using both Hymenoptera (n=4,415) and Insecta (n=1,658) Benchmarking Universal Single-Copy Orthologs (Waterhouse et al. 2018). See the Supplementary Online Material for full details of software versions and options/parameters used for each analysis.

**Figure 2.**
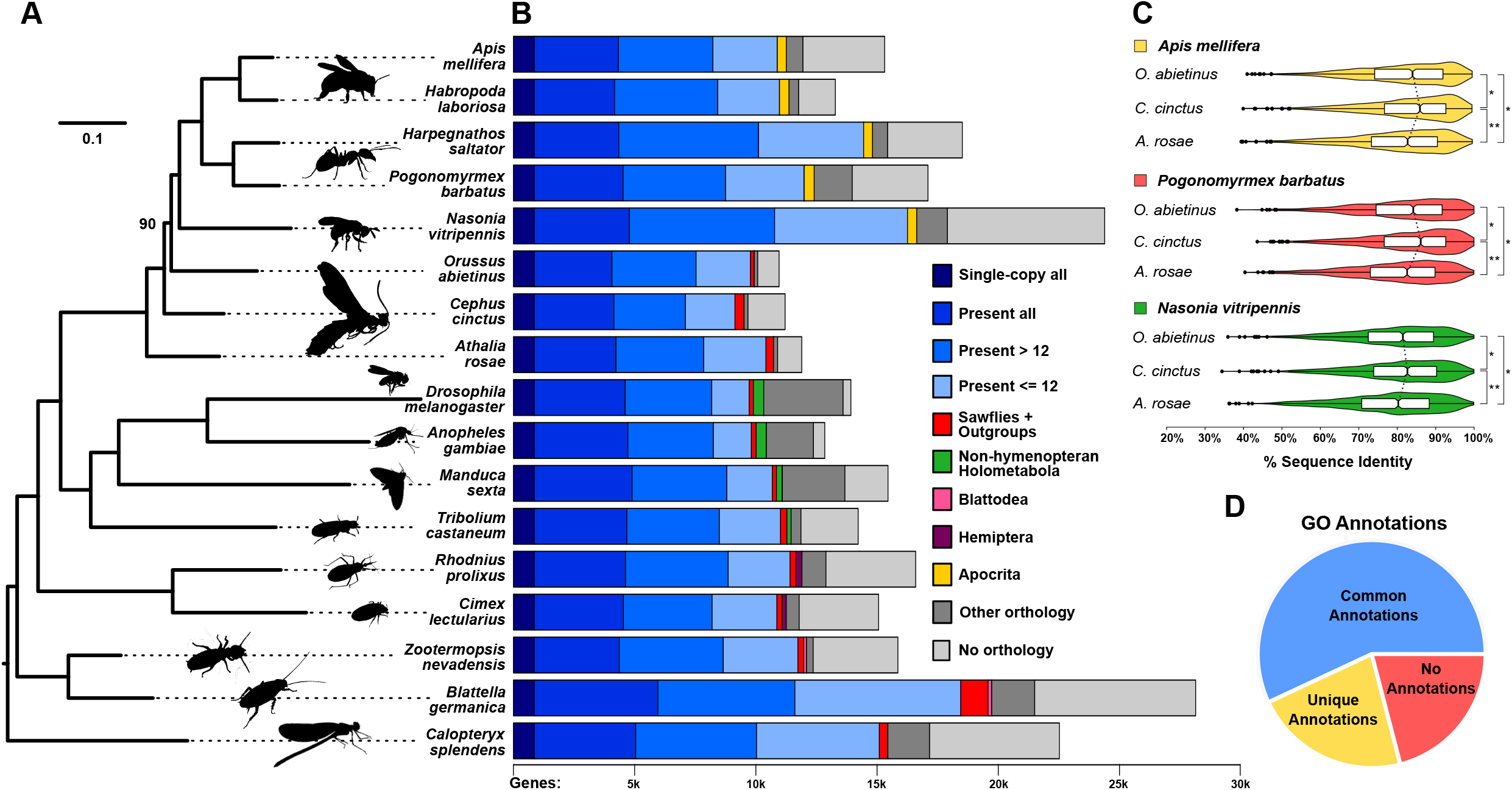
Phylogenomics, orthology, sequence identity, and functional annotation of *Cephus cinctus* genes. (A) Maximum likelihood molecular phylogeny estimated from the aligned protein sequences of 852 single-copy orthologs from 17 selected insect species, including three sawflies, *Athalia rosae, C. cinctus*, and *Orussus abietinus*, with the banded demoiselle, *Calopteryx splendens* (Odonata), as the outgroup. Branch lengths represent substitutions per site, all nodes except for the placement of *O. abietinus* (labeled) achieved 100% bootstrap support. (B) Total gene counts per species partitioned into categories from single-copy orthologs in all 17 species, present (i.e. allowing gene duplications) in all or most (>12) or with patchy species distributions (<=12), to lineage-restricted orthologs (sawflies and outgroups, non-hymenopteran Holometabola, Blattodea, Hemiptera, or Apocrita), genes with orthologs from other sequenced insect genomes, and those with currently no identifiable orthologs. (C) Distributions of percent amino acid identities of 1,035 single-copy orthologs from each of the three sawflies with *Apis mellifera, Pogonomyrmex barbatus*, and *Nasonia vitripennis*. Violin plots show the smoothed densities of each distribution, with boxplots indicating medians and upper and lower quartiles. Asterisks indicate statistically significant differences with p-values <1e-108 (**) and <1e-39 (*). (D) Functional inferences for *C. cinctus* genes from Gene Ontology (GO) annotations of orthologs from *A. mellifera, Anopheles gambiae, Drosophila melanogaster*, or *Tribolium castaneum*. Of the 7,595 *C. cinctus* genes with orthologs from any of these four species, 57% were in orthologous groups with GO terms shared amongst at least two of the four species (common annotations), 22% with GO terms from only one species (unique annotations), and 21% where no GO terms were associated with any of the orthologs (no annotations).

### Transposable and repeat element annotation

Although a repeat analysis for masking purposes is part of the NCBI gene annotation steps, we undertook a dedicated analysis using the REPET package (Quesneville et al. 2005; Flutre et al. 2011). The computational pipeline to predict and classify transposable element (TE) integrations used both *de novo*- (TEdenovo) and homology-based methods (TEannot). A summary of the REPET pipeline can be found at https://urgi.versailles.inra.fr/Tools/REPET and detailed description of the applications and methods are in the Supplementary Online Material.

### Expression Analysis

Levels of gene expression were determined by aligning the trimmed RNAseq reads from each library against the chemoreceptor and OBP transcripts from the i5k Workspace@NAL, occasionally truncated to avoid overlap with the UTRs of well-expressed neighboring genes, using the Burrows-Wheeler Aligner (BWA) (Li and Durbin 2009) and Read1 of each read pair. Samtools (Li et al. 2009) was used to sort, index, and summarize the BWA. Read counts were standardized as Read counts Per Kilobase transcript per Million mapped reads in each library (RPKM) to facilitate comparisons across the 17 samples (Table S1).

## Results

### A high quality genome assembly and annotation

The draft genome assembly consists of 10,707 contigs with a N50 of 45 kbp that were connected into 1,976 scaffolds with a N50 scaffold size of 622 kbp and a total of 162 Mbp including 3 Mbp of gaps between contigs within scaffolds. This assembly (v1) is available from the National Center for Biotechnology Information (NCBI) as BioProjects PRJNA297591 and PRJNA168335 (NCBI GCF_000341935.1). It is of a size and quality comparable to those of other Hymenoptera in GenBank (Table S2). We assessed the completeness of this assembly using BUSCO (Benchmarking Universal Single-Copy Orthologs) (Waterhouse et al. 2018), which revealed high completeness scores of 97.6-99.5%, with few duplicated (0.3-0.7%), fragmented (0.3-1.7%), or missing (0.2-0.7%) genes. These assembly completeness estimates are on a par with or marginally better than for the other assessed hymenopteran genomes (Table S4). In addition, 86-94% of the ILLUMINA RNAseq reads described below mapped to the assembly (NCBI *C. cinctus* Annotation Release 100 - https://www.ncbi.nlm.nih.gov/genome/annotation_euk/Cephus_cinctus/100/). These high levels of completeness and RNAseq read mapping indicate that this draft genome is a high quality assembly that supports effective automated and manual gene modeling, as demonstrated below.

To support gene modeling and assess gene expression across lifestages and tissues, we performed paired-end ILLUMINA RNAseq on three samples of larvae, two samples each of adult males and females, one sample each of male and female pupae, and one sample each of antennae, heads without antennae, abdomen tips, and abdomens without tips from adult males and females (Table S1 – Auxillary file 1). These libraries ranged in size from 3.5-12 Gbp. This was additionally supplemented by an available RNAseq dataset of 717,345 454-pyrosequencing reads from mixed-sex antennae (Gress et al. 2013). Gene modeling was performed by NCBI using their GNOMON pipeline, and yielded 11,210 protein-coding genes and 41 pseudogenes, as well as 1,307 non-coding RNA genes. A total of 25,937 transcripts were modeled, with 25,189 of those having support from the RNAseq and/or homology with other arthropod proteins. The mean number of transcripts per gene is 2.3 (range 1-50) and the mean number of exons per gene is 8.9 (range 1-108). The protein-coding gene set is comparable in completeness (as estimated by alignment to the REFSEQ proteins from *D. melanogaster)* with that of the eusymphytan sawfly *Athalia rosae* and the woodwasp *Orussus abietinus* (NCBI *C. cinctus* Annotation Release 100 - https://www.ncbi.nlm.nih.gov/genome/annotation_euk/Cephus_cinctus/100/). BUSCO analysis of the quality of this automated annotation in terms of expected gene content reflected the high BUSCO completeness scores for the genome assembly, identifying 98.7-99.4% complete, and few duplicated (0.4-0.7%), fragmented (0.2-0.7%), or missing (0.4-0.6%) genes. The close match of BUSCO scores for the assembly and annotations indicate that the annotation strategy was successful. These annotation completeness estimates are on a par with or marginally better than for the other assessed hymenopteran genomes (Table S4). This automated gene set therefore provides a confident base for comparative genomics, as well as subsequent manual gene annotation.

### Transposable and repeat element content

The repeat prediction pipeline identified 51,432 simple sequence repeats (SSRs) of unit length ≥ 2 that encompassed 1.53 Mbp (~1% of the assembled genome), within which hexanucleotide repeats were most abundant (Figure S1). A total of 64,215 TE fragments totaling 20.8 Mbp (~13% of the assembled genome) were resolved into 57,948 predicted copies using the Long_join method to connect disrupted portions of the same integration. These predictions are available as a track at the i5k Workspace@NAL browser for this genome (https://i5k.nal.usda.gov/cephus-cinctus). The PASTEC classification of predicted repeat elements from the REPET TEdenovo pipeline identified 840 unique elements placed into six Class-I families and five Class-II families (Table S5). Overall, although the copy number of predicted elements are approximately equal between Class-I (n = 21,429) and Class-II (n = 21,473) elements, the latter occupies ~2-fold greater portion of the genome (Class-I: 5.3 Mbp vs. Class-II: 10.2 Mbp). Among the Class-I retroelements, long-interspersed nuclear elements (LINEs) are the most abundant, but long terminal repeat (LTR) elements occupy a greater proportion of the genome. Incomplete copies of Class-II elements across all families are most abundant, including among the non-autonomous short interspersed nuclear element (SINE) and terminal-repeat retroelements in miniature (TRIM) elements (Table S5). Inverted repeat (TIR) family members are most abundant and encompass the greatest proportion of the genome among Class-II DNA elements. The miniature inverted repeat transposable elements (MITEs) are the second most abundant family, but likely due to being the only fully non-autonomous family of Class-II elements they occupy a smaller proportion of the genome than TIR or *Maverick* elements.

### Phylogenomics and protein orthology

The molecular species phylogeny (Figure 2A), estimated from concatenated protein sequence alignments of single-copy orthologs, supports the current view of sawfly radiations early during the evolution of the Hymenoptera (Peters et al. 2017), with Cephidae placed between the more basal Tenthredinidae (represented by the turnip sawfly, *A. rosae*) and the more derived Orussidae (represented by the parasitic wood wasp, *O. abietinus*). Orthology delineation across insects identified orthologs for 86.4% of *C. cinctus* genes, almost three quarters of which have orthologs in all or most of seven other hymenopterans and nine representative species from six other insect orders (Figure 2B). The two sawflies and the woodwasp have similarly low total gene counts and show similar proportions of genes in each orthology category. In contrast, the other representative hymenopterans have generally higher total gene counts including a fraction of seemingly apocritan-specific orthologs.

The small fraction of sawfly genes with orthologs in outgroup species, but not in other Hymenoptera, highlight potential gene losses in the apocritan ancestor. These include orthologs of *D. melanogaster* genes *TrpA1* (*Transient receptor potential cation channel A1*) involved in responses to heat and noxious chemicals (Luo et al. 2017), and the G-protein-couple receptor *Lgr1* (*Leucine-rich repeat-containing G protein-coupled receptor 1*) involved in development (Vandersmissen et al. 2014). This fraction also includes a gene encoding a CPCFC family cuticular protein, previously described by Vannini et al. (2015) as being present in *C. cinctus* but missing from apocritan species. Further examples of putative losses in the Apocrita ancestor, from our detailed analysis of chemoreceptor gene families, are elaborated below.

Examining ortholog sequence conservation between sawfly and Apocrita species showed that *C. cinctus* proteins had significantly higher amino acid identities (Wilcoxon signed rank tests, p<1e-39) with the Apocrita than either *A. rosae* or *O. abietinus* (Figure 2C). This apparently slower rate of sequence divergence in *C. cinctus* may at least partially explain the uncertainty of the placement of *O. abietinus* in the species phylogeny (Figure 2A), which was alternatively placed as sister to *C. cinctus* in 10% of bootstrap samples. With little detailed functional characterization of *C. cinctus* genes to date, putative functions can instead be inferred by identifying orthologs from well-studied insects such as the honey bee, malaria mosquito, fruit fly, or flour beetle, to tentatively link Gene Ontology terms to the majority of *C. cinctus* genes (Figure 2D).

### microRNAs

A total of 36 putative high confidence miRNAs from 22 different miRNA families were identified. Of these, 24 were located on the sense strand while 12 were on the antisense strand (Table 1).

**Table 1.** miRNA ID and number of miRNA annotations.

| **mirnaID** | **on**<br>**SENSE** | **on**<br>**ANTISENSE** | **total** |
| --- | --- | --- | --- |
| miR-10-5p | 2 | 0 | 2 |
| miR-137-3p | 2 | 0 | 2 |
| miR-1-3p | 1 | 0 | 1 |
| miR-14-3p | 0 | 2 | 2 |
| miR-1-5p | 1 | 0 | 1 |
| miR-184-3p | 1 | 0 | 1 |
| miR-190-5p | 0 | 2 | 2 |
| miR-210-3p | 1 | 0 | 1 |
| miR-210-5p | 1 | 0 | 1 |
| miR-275-3p | 0 | 1 | 1 |
| miR-277 | 0 | 1 | 1 |
| miR-279-3p | 0 | 1 | 1 |
| miR-2796-3p | 1 | 0 | 1 |
| miR-281-3p | 3 | 0 | 3 |
| miR-281-5p | 1 | 0 | 1 |
| miR-2a-3p | 0 | 1 | 1 |
| miR-315-5p | 0 | 1 | 1 |
| miR-71-5p | 0 | 1 | 1 |
| miR-8-3p | 2 | 0 | 2 |
| miR-8-5p | 1 | 0 | 1 |
| miR-87a-3p | 0 | 1 | 1 |
| miR-927a-5p | 1 | 0 | 1 |
| miR-929-5p | 1 | 0 | 1 |
| miR-92b | 1 | 0 | 1 |
| miR-92b-3p | 3 | 0 | 3 |
| miR-981-3p | 1 | 0 | 1 |
| miR-iab-4-5p | 0 | 1 | 1 |
| **SUM** | **24** | **12** | **36** |

### The C. cinctus chemosensory gene repertoire

Insects depend on members of three major gene families for most of the sensitivity and specificity of their senses of smell and taste (Leal 2013; Joseph and Carlson 2015). The first two are the Odorant Receptor (OR) and Gustatory Receptor (GR) families, which together form the insect chemoreceptor superfamily (Robertson et al. 2003) of seven-transmembrane ligand-gated ion channels. The third family are the Ionotropic Receptors (IRs) that are evolutionarily unrelated to the insect chemoreceptor superfamily, being a variant lineage of the ionotropic glutamate receptor superfamily (Benton et al. 2009; Ryzt et al. 2013; van Giesen and Garrity 2017; Rimal and Lee 2018). Our manual annotation focused on these three gene families, as well as the Odorant Binding Protein (OBP) family of secreted small globular proteins, resulting in high-quality gene models from which to build robust inferences of their evolutionary histories.

The OR family consists of a highly conserved Odorant receptor co-receptor (Orco) gene that has 1:1 orthologs throughout the pterygote insects (Ioannidis et al. 2017) and forms dimers with each of the many “specific” ORs (Benton et al. 2006). In *Cephus* there are 72 OR genes including the expected single highly conserved *Orco* (Table S6), a total considerably smaller than the large OR families of *A. mellifera* (176 proteins – Robertson and Wanner 2006), *B. terrestris* (166 – Sadd et al. 2015), *P. barbatus* (400 – Smith et al. 2011), and *N. vitripennis* (301 – Robertson et al. 2011). Phylogenetic analysis using protein sequences from these species, employed because they have undergone similarly intensive manual annotation, reveals that the *Cephus* ORs are largely present as a single gene, or sometimes small clusters of genes, at the base of numerous small clades and sometimes highly-expanded subfamilies in these other hymenopterans (Figure S2). Most remarkably, a single gene (*CcinOr69*) is at the base of the extremely large 9-exon subfamily, which is particularly highly expanded in ants and mediates perception of their cuticular hydrocarbons of crucial importance to social interactions (Smith et al. 2011; Pask et al. 2017). Similarly, a small species-specific clade consisting of *CcinOr1-9* is at the base of the large-tandem-array subfamily of 61 genes in *A. mellifera* (Robertson and Wanner 2006) that includes a receptor for the primary queen pheromone substance, *AmelOr11* (Wanner et al. 2007), a tandem array that has persisted throughout hymenopteran evolution. In addition to this small Cephus-specific clade of tandem-array subfamily members, there are three more Cephus-specific clades of 4, 8, and 8 genes, which are extremely small compared with the large gene subfamily expansions observed in the apocritans. *Cephus* appears to have lost a few OR lineages present in the other hymenopterans, and has retained several lineages that are missing from the Apocrita but are commonly still present in other symphytans and/or the wood wasp. This analysis suggests that ancestral hymenopterans had approximately 40 OR lineages.

The GR family is a highly diverse family of primarily taste receptors, but also some olfactory receptors such as the carbon dioxide receptors of flies and moths, including receptors for sugars and a vast array of “bitter” compounds, as well as light and heat. It is an ancient family within the Metazoa, dating back to basal animals (Saina et al. 2015; Robertson 2015; Eyun et al. 2017), but was lost from vertebrates, which instead employ primarily G-protein-coupled receptors for both olfaction and taste. We identified a total of 35 GR genes in *C. cinctus* (Table S7), an intermediate number compared with the other species with 13 in *A. mellifera* (Robertson and Wanner 2006), 25 in *B. terrestris* (Sadd et al. 2015), 75 in *P. barbatus* (Smith et al. 2011), and 47 in *N. vitripennis* (Robertson et al. 2011). In addition to single orthologs of the two sugar receptors known for hymenopterans *(CcinGr1/2)* and a single candidate fructose receptor *(CcinGr3), Cephus* has two clear orthologs of the carbon dioxide receptors of other endopterygote insects *(CcinGr24/25)*, although the third carbon dioxide receptor GR lineage is missing (Figure 3). In addition, there are two more genes clearly related to the carbon dioxide receptor clade *(CcinGr26/27)*, which are part of an expanded subfamily in exopterygote insects from which the carbon dioxide receptors evolved (Ioannidis et al. 2017). The remaining GRs, like the ORs, are placed in the phylogeny as single genes or small species-specific clades at the base of most GR lineages and subfamilies, including several large expansions in ants and wasps. There are again several highly divergent GR lineages in *Cephus* that are commonly present in other symphytans and the woodwasp, but have been lost from the more derived Hymenoptera, specifically CcinGr16-20, 21, 23, 28, 29-32, 33-36 (Figure 3). Unlike for the ORs, there are no obvious losses of entire GR lineages from *Cephus*. These comparisons lead to an estimate of approximately 30 GR lineages in ancestral hymenopterans, maintained in *Cephus* but with some lineages lost from, and others greatly expanded in, the Apocrita.

**Figure 3.**
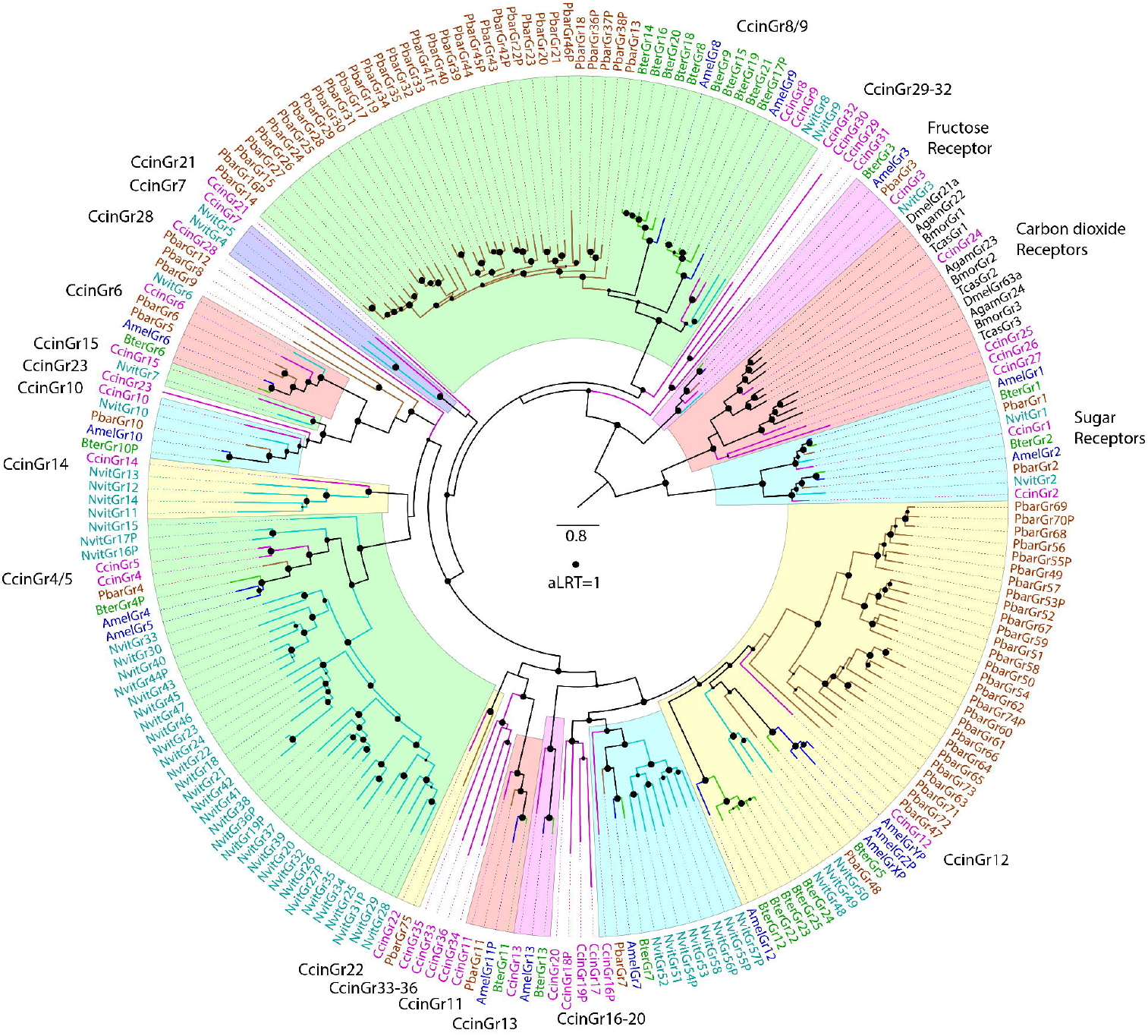
Phylogenetic relationships of the Gustatory Receptor family in Hymenoptera. The tree was rooted by declaring the sugar and carbon dioxide receptor lineages as the outgroup, based on their basal location within insect GRs in an analysis of the entire GR family in insects and other animals (Robertson 2015). GR names and the branches leading to them are colored by species, purple for *C. cinctus*, cyan for *N. vitripennis*, brown for *P. barbatus*, green for *B. terrestris*, blue for *A. mellifera*, and black for other species. The CcinGr lineages are indicated outside the circle of names, and major lineages with CcinGr representatives are highlighted in colored segments. The scale bar indicates substitutions per site, and the filled circle indicates approximate likelihood ratio test (aLRT) support of 1.

The IRs have an extracellular ligand-binding domain supported by three transmembrane domains, and also function as ligand-gated ion channels. They are involved in both olfaction, primarily being expressed in olfactory sensory neurons in coeloconic sensilla that do not express ORs and sensing acids and amines, and gustation with large subfamilies expressed in gustatory organs, at least in *Drosophila melanogaster* (Rytz et al. 2013; Koh et al. 2014). A few are also involved in sensing temperature and humidity (Knecht et al. 2017; van Giesen and Garrity 2017; Rimal and Lee 2018). The IR family in *C. cinctus* is made up of 49 genes (Table S8), roughly double the number of *A. mellifera* (21 – Croset et al. 2010), *B. terrestris* (22 – Sadd et al. 2015), and *P. barbatus* (27 – Smith et al. 2011 and supplement), but considerably smaller than *N. vitripennis* for which our manual annotation efforts increased the gene count from just 10 in Croset et al. (2010) to 153 genes, albeit 54 (35%) are pseudogenes (supplement). *Cephus* has the expected single orthologs of the highly conserved *Ir8a, 25a*, and *76b* genes, which encode proteins that function as co-receptors with other IRs (Rytz et al. 2013). It also has single orthologs for three of the four genes involved in thermo- and hygro-sensation in *D. melanogaster (Ir21a, 68a*, and *93a)*, but *Ir40a* was lost at the base of the Hymenoptera. Ir40a cooperates with the other proteins to sense humidity in *D. melanogaster* (Knecht et al. 2017), so it would be interesting to discover how this gene loss, and idiosyncratic losses of *Ir21a* and *Ir68a* in various Hymenoptera, affect their abilities to perceive temperature and humidity. *Cephus* has only a single relative of the *DmelIr75a-c* genes that encode olfactory receptors for various acids (Prieto-Godino et al. 2017), a lineage that is expanded in wasps, but of comparable size to flies in ants and bees. There is a single hymenopteran set of orthologs for the *DmelIr41a* lineage of olfactory receptors that is commonly greatly expanded in other insects (Robertson et al. 2018). Like the ORs and GRs, *Cephus* has generally single representatives of other hymenopteran IR lineages, but also two Cephus-specific clades, one of several ancient gene lineages that appear to have been lost from Apocrita, and one of very recent gene duplications (Figure S3). These clades, along with a few bee and ant proteins and the majority of the *Nasonia* proteins, are related to the DmelIr7a-f/11a proteins and the large DmelIr20a clade implicated in gustation (Rytz et al. 2013; Koh et al. 2014). This analysis suggests that ancestral hymenopterans had approximately 34 IR gene lineages. Thus *Cephus* has the expected complement of conserved hymenopteran IRs, except for the *Ir40a* and *Ir41a* lineages, as well as both ancient lineages and very recent species-specific clades, the latter of which, like those in the OR and GR families, might be involved in colonization of new grass species including wheat.

Finally, OBPs are small globular proteins commonly secreted by support cells at the base of chemosensory sensilla and thought to be involved in transport of commonly hydrophobic odorants across the chemosensillar lymph (Pelosi et al. 2014), although not all OBPs are expressed in chemosensory tissues (Foret and Maleszka 2006; Pelosi et al. 2018). Furthermore, the only experimentally demonstrated biological role for an OBP is that of quenching the signal of an odorant (Larter et al. 2016). Like the chemoreceptor families, hymenopterans have a wide range of numbers of OBPs, from 18 in the fire ant *Solenopsis invicta* (Gotzek et al. 2011) and 21 in *A. mellifera* (Foret and Maleszka 2006) to 90 in *N. vitripennis* (Vieira et al. 2012). We identified 15 OBP genes and their encoded proteins in *C. cinctus* (Table S9), which like the chemoreceptors are frequently at the base of a clade of single orthologs or expansions in these other hymenopterans (Figure S4).

### Expression levels of chemoreceptors and OBPs

We examined the levels of expression of the chemoreceptors and OBPs in our 17 diverse RNAseq datasets from larvae, pupae, adults, and adult body parts (TableS1 – Auxiliary file 1), using the manually annotated transcripts, truncated if necessary to avoid conflation with overlapping untranslated regions (UTRs) of highly-expressed neighboring genes. The complete results are presented in Tables S10-13 and Figures S5-8 (Auxiliary file 2), with selected genes shown in Figures 4-7. For the ORs, *Orco* has the expected high levels of expression in male and female antennae, roughly twice the level seen in entire adult bodies or pupae, where chemoreceptor expression is initiated. As expected, heads without antennae, abdomen tips, and abdomens without tips had low levels of expression of *Orco*. Larvae also have low levels, perhaps because larvae only have a few olfactory sensilla given their feeding mode within wheat stems. These expected results for *Orco* provide confidence that our other results are reliable.

Most OR genes have appropriately low expression, primarily in antennae (Figure S5), represented by *Or5* in Figure 4, while some of the divergent genes like *Or60* and *Or72* are barely expressed. There are a few instances of consistent sex bias. For example, *Or1* is consistently expressed ~4X higher in males than females, while *Or30-32* exhibit 5-7X higher expression in males, although these three might be conflated as they are very similar genes in a tandem array (represented by *Or32* in Figure 4). All four are amongst the mostly highly expressed Ors, a result confirmed by prior quantitative PCR that also found *Or32* expression 15X higher in male antennae (Gress et al. 2013; Table S14), making these candidate receptors for female pheromone. The expression of *Ors 18, 63, 65* and *68* was higher in female antennae, in this study and in prior quantitative PCR experiments (5-15X female bias, Gress et al. 2013; Table S14), making these candidate receptors for host plant volatiles or male-produced pheromones.

**Figure 4.**
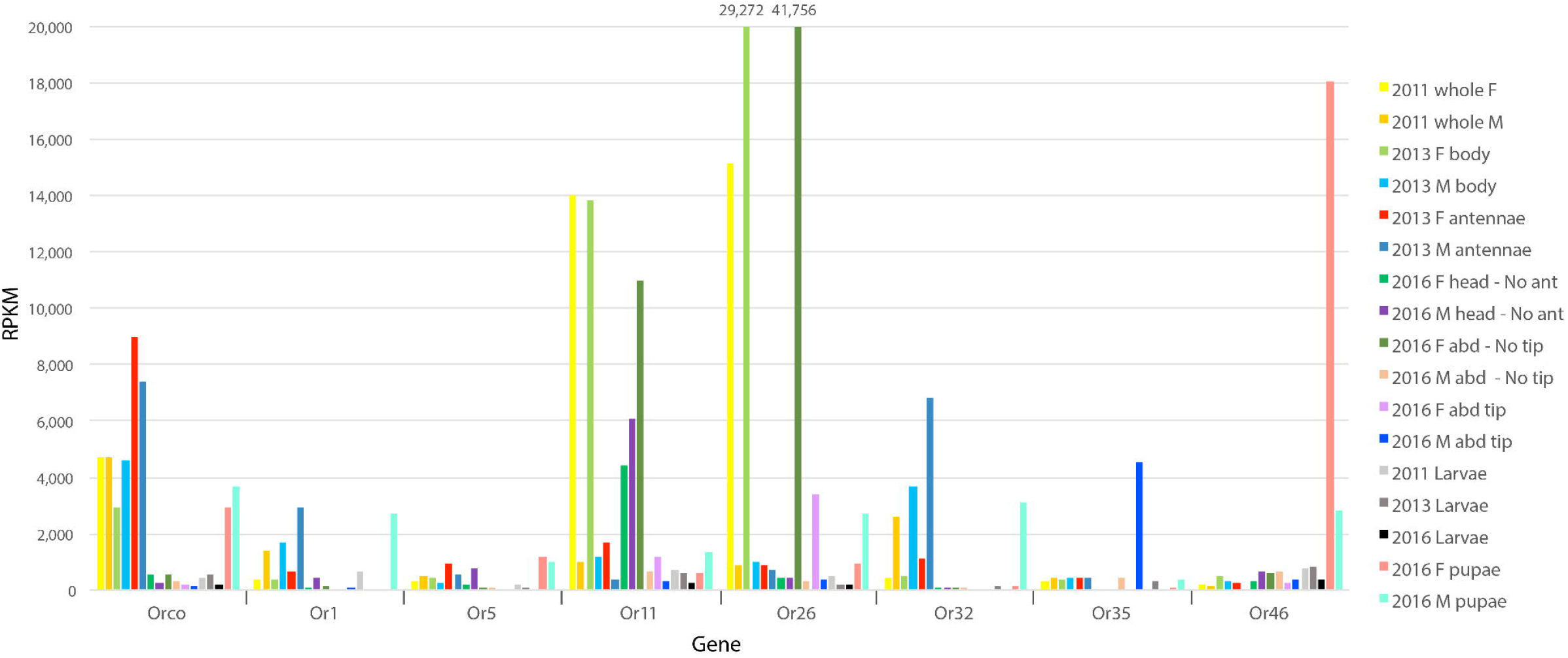
Expression levels of selected ORs. Expression levels are shown as Reads Per Kilobase transcript per Million reads (RPKM) in each library for eight of the 73 OR family genes and the 17 RNAseq libraries (detailed in Table S1) that sampled different life stages and tissues of male and female sawflies collected from 2011 to 2016. Abbreviations: abd, abdomen; ant, antennae; F, female; M, male.

Some ORs exhibit unusual expression patterns, for example, the tandemly-arrayed *Or11-13* are expressed at levels comparable to *Orco* in heads without antennae, and *Or11* is also extremely highly expressed in female bodies, specifically abdomens. It would be interesting to see if the relatives of this small clade of *CcinOr11-14* in other Hymenoptera (*NvitOr11-33, PbarOr70-75, BterOr87-95*, and *AmelOr114/5*) are similarly expressed outside the obvious olfactory organs. A few other unusual results are high expression of *Or35* and *Or65* in male abdominal tips, and of *Or46* in pupae. An enigmatic result is the extraordinarily high expression of *Or26* in females, specifically their abdomens. The 3’ UTR of this gene overlaps that of a neighboring highly-expressed DNA polymerase gene, hence the analyzed transcript was shortened at the 3’ end, so this result has to be treated with caution, nevertheless the reads that map to *Or26* are mostly correctly spliced, implying they are from this locus rather than the 3’ UTR of the polymerase.

Amongst the GRs, as expected the sugar receptors *Gr1/2* are expressed primarily in heads at levels comparable to or higher than *Orco*, however *Gr2* is also expressed in abdomens and abdominal tips, and unusually highly in female abdomens. The candidate fructose receptor *Gr3* is the mostly highly expressed GR, and as expected from the expression of its *Drosophila* ortholog *DmelGr43a* (Miyamoto et al. 2012), primarily in heads but also in abdomens, larvae, and pupae, and appropriately low in antennae. Most of the other GRs show the expected pattern of low levels of expression primarily in heads (Figure S6), rather than antennae, a pattern exemplified by *Gr36* (Figure 5). Exceptions include *Gr12*, which is consistently highly expressed in females, primarily in abdomens. The two candidate carbon dioxide receptors, *Gr24/25*, are also not well expressed in antennae, but rather in heads, and *Gr24* is unusually highly expressed in pupae. The related carbon-dioxide receptor subfamily gene *Gr26* shows a similar pattern (unfortunately *Gr27* overlaps the 3’ UTR of a highly-expressed gene, so could not be evaluated).

**Figure 5.**
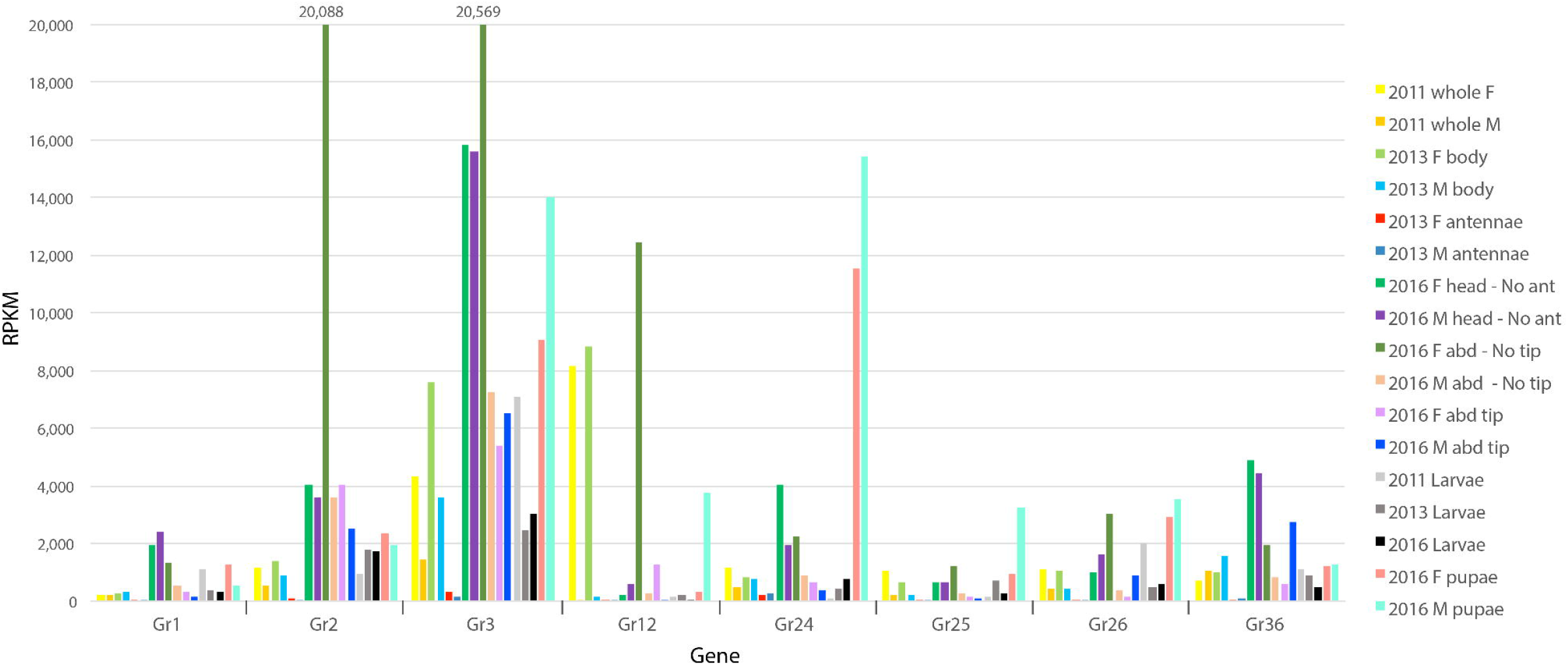
Expression levels of selected GRs. Expression levels in RPKM for eight of the 36 GR family genes. Other details as for Figure 4.

Amongst the IRs, *Ir25a* and *76b*, two co-receptors, are the most highly expressed, primarily in heads, as are a few others like *Ir41a*, however *Ir76b* is extremely highly expressed in male abdominal tips, a clear result with almost entirely spliced reads. The third co-receptor, *Ir8a*, is expressed at a relatively low level. *Ir93a* is the most highly expressed of the three genes implicated in perception of temperature and humidity, with *Ir68a* the lowest of these three. It is noteworthy that these are the more conserved receptors whose orthologs in *D. melanogaster* are primarily expressed in antennae (Rytz et al. 2013). Amongst the more divergent IRs, the most highly expressed are *Ir108* and *Ir120*_¦_, again primarily in heads, while *Ir140* is shown as an example of the generally low expression of the IRs with likely gustatory roles (Figure 6).

**Figure 6.**
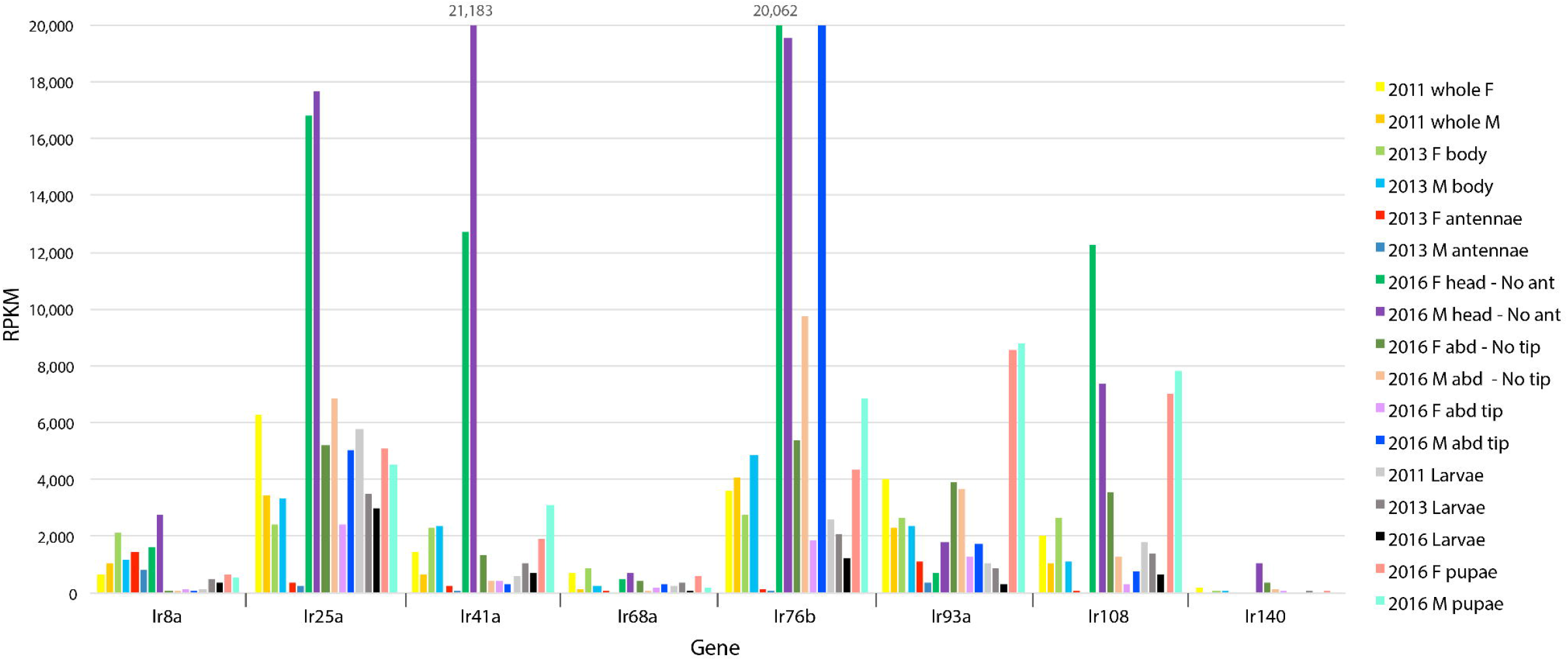
Expression levels of selected IRs. Expression levels in RPKM for eight of the 49 IR family genes. Other details as for Figure 4.

Finally, as expected the OBPs commonly exhibited far higher levels of expression than the chemoreceptors (Figure S8). Perhaps surprisingly, only *Obp1, Obp4, Obp6, Obp11* and *Obp12* are well expressed in antennae (and male abdominal tips for *Obp12)* (represented by *Obp6* and *Obp12* in Figure 7). *Obp2* and *Obp9* are primarily expressed in heads (and larvae for *Obp9)*, while *Obp3, Obp5, Obp8* and *Obp15* have unusually low levels of expression for OBPs, and mostly in larvae. *Obp7* and *Obp10* are primarily expressed in male abdomens, especially the tips, while *Obp13* is almost exclusively expressed in larvae. Finally, *Obp14* is extraordinarily highly expressed in multiple samples, most highly in female abdominal tips. The observation of high expression of several OBPs in abdominal tips implies a non-olfactory role for these proteins.

**Figure 7.**
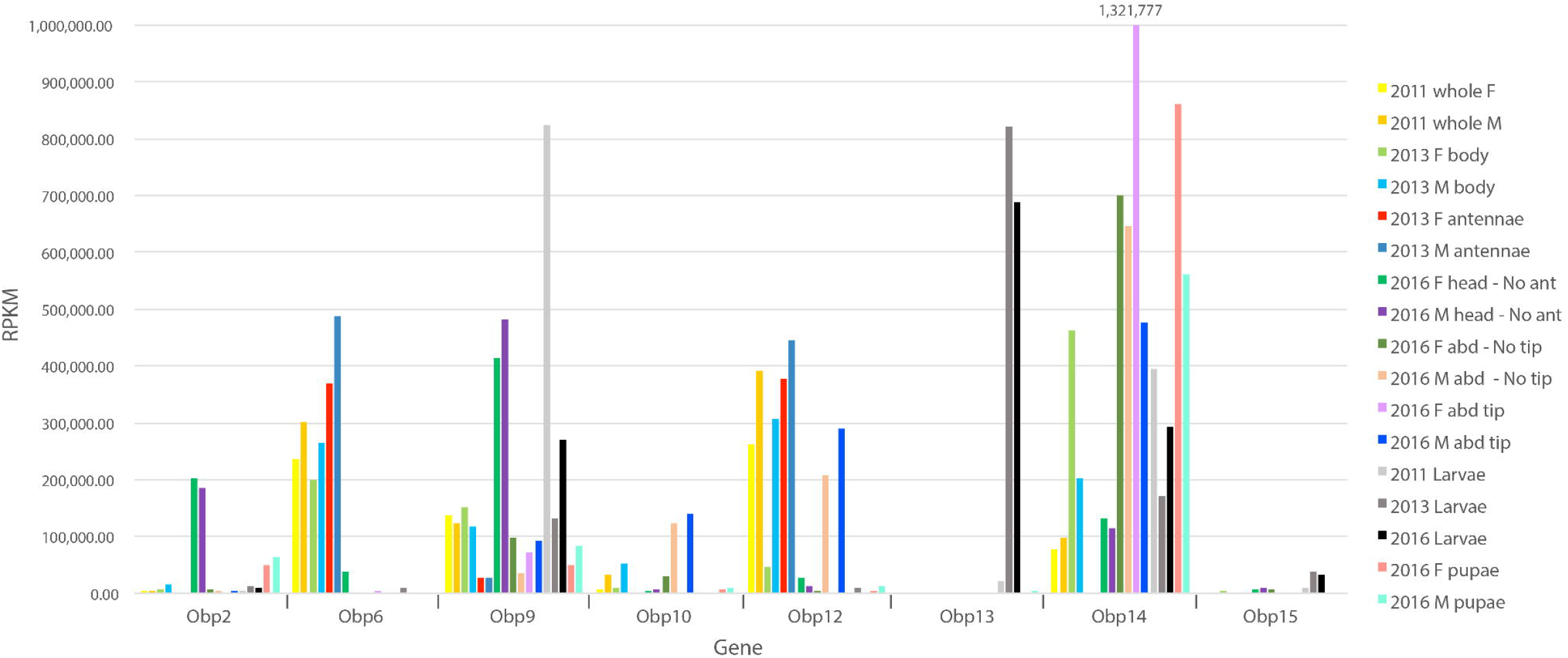
Expression levels of selected OBPs. Expression levels RPKM for eight of the 15 OR family genes. Other details as for Figure 4.

## Discussion

Genome sequencing offers the opportunity to explore the putative genomic or genetic basis of organismal biology that may be widely shared among species or highly lineage-specific. The informed interpretation of observed patterns of conservation and divergence requires a robust understanding of the evolutionary relationships among the considered species, which in the case of sawflies has remained unclear until relatively recently. While the separation of Cephidae from the other symphytan sawflies had been proposed for some time, it was only with the phylogenomics study of Peters et al. (2017), based on protein sequences deduced from transcriptomes, that this placement was put on more solid footing. Our genome-based phylogenomics analysis cements the case for placement of the Cephidae as a secondary radiation after the Eusymphyta, and with considerable confidence agrees with Peters et al. (2017) in the branching of the Cephidae before the orussid woodwasps. The apparently constrained rates of molecular evolution of *Cephus* genes relative to other sawflies and woodwasps, resulting in higher amino acid identities with their Apocrita orthologs, may partially explain previous difficulties in attempting to resolve these early hymenopteran radiations.

The availability of many other hymenopteran and other insect genomes means that the vast majority (86%) of *Cephus* proteins can be confidently assessed as having orthologs in other species, despite the deep branching of this lineage. Functional information from orthologs in *D. melanogaster* and other insects allows assignment of putative biological roles for more than half of the annotated *C. cinctus* protein-coding genes. These associations reflect the ever-increasing knowledge of insect molecular biology and provide a strong foundation for future functional work on this sawfly.

Our repeat analysis from this 162 Mbp genome assembly revealed that 1% consists of simple sequence repeats, amongst which are just 27 copies of the TTAGG telomeric repeat found to constitute the telomeres of the honey bee (Robertson and Gordon 2006). None of these TTAGG repeats are obviously at the end of long scaffolds where they might constitute telomeres, indicating that despite the high quality of our assembly, telomeric regions, and surely centromeric regions, remain unassembled, as is the case for most draft genome sequence assemblies. A further 13% of the assembly consists of transposons, partitioned roughly equally between Class-I and Class-II elements. This TE profile is in contrast to that seen in the honey bee where simple repeats make up 4% of the 250Mbp genome assembly and transposons comprise only 5% (Elsik et al. 2014). We still have only a rudimentary understanding of how genomes come to have such disparate transposon compositions.

Our analysis of the three major chemoreceptor families and the OBPs reveals *Cephus* genes at the base of most apocritan gene lineages, many as single-copy orthologs but also several with remarkably large expansions in wasps, ants, and/or bees. While only a few gene lineages appear to have been lost from *Cephus*, several are absent in the apocritans, most prominently the entire carbon dioxide receptor subfamily. This pinpoints the loss to the apocritan ancestor, and leaves open the question of how extant apocritans are able to sense carbon dioxide concentrations. *Cephus* nevertheless does have a few small species-specific expansions of chemoreceptors that might mediate some relatively recent adaptations to new grasses including wheat, which could be important for future analysis of phylogeography of this recently adapted crop pest (Lesieur et al. 2016). Our identification of candidate pheromone and host volatile receptors, as well as gustatory receptors expressed on the ovipositor will inform future studies of behavioral and chemical ecology (Varella et al. 2017). Examination of gene expression levels reveals generally expected results, such as high expression of *Orco* in antennae and the sugar and fructose GRs and the IR co-receptors in heads, with low expression of most other receptors. Several exceptions are noted including sex-biased and tissue-specific expression. This complete cataloging of these chemosensory gene families now makes detailed functional studies of their involvement in the ecology of this sawfly feasible.

In summary, this high-quality draft genome assembly and automated gene annotation will facilitate further molecular biological studies on this sawfly, and offers opportunities to identify genes for engineering of sawfly-resistant wheat strains with RNAi constructs. It also augments the diversity of hymenopterans with sequenced genomes, particularly for the poorly sampled sawflies (Branstetter et al. 2018), making this insect order second only to Diptera for the number of species with genome sequences.

## Acknowledgements

We thank Alvaro Hernandez and the W. M. Keck Center for Comparative and Functional Genomics at the University of Illinois at Urbana-Champaign for genomic and RNA library construction and sequencing; Daniel Ence and Mark Yandell (University of Utah) for annotation of an earlier version of the genome assembly; Terence Murphy for assistance with the NCBI annotation; Monica Poelchau and Chris Childers for assistance with the i5k Workspace@NAL browser and official gene set v1.1; Chris Elsik for an Apollo browser at NasoniaBase; Masatsugu Hatakeyama, Bernhard Misof, Oliver Niehuis, and Jan Philip Oeyen for granting pre-publication access to gene annotations of *Athalia rosae* and *Orussus abietinus*; and Megan Hofland and Norma Irish for preparing Figure 1. This work was supported by funding from the United States Department of Agriculture [grant number AG2008-35302-188815 to HMR and KWW], Swiss National Science Foundation [grant number PP00P3_170664 to RMW], the Winifred-Asbjornson Plant Science Endowment to HB, the Montana Wheat and Barley Committee and the Montana Grains Foundation to DKW and KWW, who also received funds supporting this research from the AES allocation in HB 645 of the 61^st^ Legislature of the State of Montana, and a contribution from Gene Robinson (University of Illinois at Urbana-Champaign). Part of this research was the result of a joint contribution from the United States Department of Agriculture (USDA), Agricultural Research Service (ARS) (CRIS Project 5030-22000-018-00D), and the Iowa Agriculture and Home Economics Experiment Station, Ames, IA (Project 3543). USDA is an equal employment opportunity provider. This article reports the results of research only.

